# Modulation of human kinase activity through direct interaction with SARS-CoV-2 proteins

**DOI:** 10.1101/2023.11.27.568816

**Authors:** Bettina Halwachs, Christina S. Moesslacher, Johanna M. Kohlmayr, Sandra Fasching, Sarah Masser, Sébastien A. Choteau, Andreas Zanzoni, Natalia Kunowska, Ulrich Stelzl

## Abstract

The dysregulation of cellular signaling upon SARS-CoV-2 infection is mediated via direct protein interactions, with the human protein kinases constituting the major impact nodes in the signaling networks. Here, we employed a targeted yeast two-hybrid matrix approach to identify direct SARS-CoV-2 protein interactions with an extensive set of human kinases. We discovered 51 interactions involving 14 SARS-CoV-2 proteins and 29 human kinases, including many of the CAMK and CMGC kinase family members, as well as non-receptor tyrosine kinases. By integrating the interactions identified in our screen with transcriptomics and phospho-proteomics data, we revealed connections between SARS-CoV-2 protein interactions, kinase activity changes, and the cellular phospho-response to infection and identified altered activity patterns in infected cells for AURKB, CDK2, CDK4, CDK7, ABL2, PIM2, PLK1, NEK2, TRIB3, RIPK2, MAPK13, and MAPK14. Finally, we demonstrated direct inhibition of the FER human tyrosine kinase by the SARS-CoV-2 auxiliary protein ORF6, hinting at pressures underlying ORF6 changes observed in recent SARS-CoV-2 strains. Our study expands the SARS-CoV-2 – host interaction knowledge, illuminating the critical role of dysregulated kinase signaling during SARS-CoV-2 infection.

## Introduction

Severe acute respiratory syndrome coronavirus 2 (SARS-CoV-2) associated with human respiratory disease was identified in December 2019 in China [1] and quickly spread across the world causing a lethal pandemic with unprecedented impact on all areas of human life. In the research community, scientists of all disciplines launched projects to help understand the disease in all aspects, prevent the spread, ease its socio-economic impacts, and develop a vaccine and a cure. This led to SARS-CoV-2 infection being one of the best studied diseases, with many comprehensive data sets associated with it.

In particular, studies of interactions between SARS-CoV-2 and human proteins provide links to biological pathways and cellular processes impacted by the infection. These data can in turn guide the attempts to find cellular targets for pharmacological intervention in order to reverse these changes [2]. Multiple large, high-quality SARS-CoV-2 – human protein interaction data sets have been generated. The current data body stems mainly from affinity purification-mass spectrometry (AP-MS) approaches [3–7], which preferentially capture stable protein complexes. However, the data also include two more recent, larger Y2H interaction studies that report direct protein-protein host-viral interactions [7, 8].

In parallel, wide host phospho-proteome responses to infection have been measured [4, 9–13]. More than a half of the SARS-CoV-2 proteins in human cells are phosphorylated at multiple sites. Moreover, cellular host proteins also undergo extensive changes in post-translation modifications, suggesting functional dysregulation of responsible cellular enzymes attributed to the infection. However, there are hardly any clear molecular or mechanistic connections between the protein interactions, the phospho-responses and altered human kinase activities. Only in very rare cases, such as ORF9B-MARK, a direct kinase – SARS-CoV-2 protein interaction could be demonstrated [14].

From the around 120 potential interacting SARS-CoV-2 - kinase pairs that can be inferred from the protein complexes identified across the major AP-MS studies [3–7], most interactions are likely to be indirect, as the fraction of direct binary interactions in AP-MS data is estimated to be around 20% [15, 16]. In general, kinase-protein interactions are difficult to detect with biochemical purification approaches such as AP-MS [17], as they tend to be more dynamic and less abundant than other protein complexes. Such interactions are better captured by the yeast two-hybrid (Y2H) analyses, which have shown to be successful in establishing transient signaling interactions [18]. Two large scale studies made use of the Y2H system to generate a SARS-CoV-2-human protein-protein interactome network [7, 8]. These studies aimed to provide high quality binary SARS-CoV-2 human interactions, prioritizing druggable targets and employing network-based drug repurposing strategies. However, these type of proteome scale studies suffer from a relatively high false negative rate [19], missing out many protein interactions. Consequently, the two Y2H data sets [7, 8] have only 11 interactions in common. Moreover, while Zhou et.al report five and Kim et al. three interactions involving protein kinases, none of these are in the overlap. Taking the union of major AP-MS and Y2H data sets into account, no SARS-CoV-2 human kinase interaction pair is common across three or more studies while only five pairs have been identified in two studies each: M-COQ8B, M-PRKDC, nsp9-NEK9, ORF3a-EPHA2, ORF7a-ATR and ORF9B-MARKs. Despite of large infection associated changes of the human phospho-proteome, to date human kinase - SARS-CoV-2 interactions still remain understudied.

Here, we use a targeted Y2H matrix approach to screen for direct protein interactions between a large set of human kinases and SARS-CoV-2 proteins. We report 29 human kinase – SARS-CoV-2 protein interaction pairs and integrate the information on human-kinase interactions with the available transcriptomics and phospho-proteomics data. As a result, we connect the phospho-response seen upon SARS-CoV-2 infection in human cells and direct interactions of SARS-CoV-2 proteins with human kinases. Finally, as an example of a downstream effect, we show that SARS-CoV-2 ORF6 can inhibit the tyrosine kinase activity of its interaction partner FER.

## Results and Discussion

We sought to address the dearth of information regarding direct physical contacts between human kinases and SARS-CoV-2 proteins by mapping SARS-CoV-2 – human kinases interactions through systematic Y2H analyses. We first established a full SARS-CoV-2 clone set from resources shared within the research community [3, 20]. We then cloned the 29 SARS-CoV-2 ORF sequences in four Y2H vectors, creating two LexA-DNA-binding (bait) and two Gal4-activation domain constructs (prey), each encoding for either a N-terminal or a C-terminal domain fusion protein. To verify the utility and quality of this clone set, we used it to test intra-SARS-CoV-2 protein interactions (**Suppl. Figure 1**). While some of the proteins (ORFs 3a, 3b, 6, 7a, 7b, 8, 9a, 9b, and 10) did not show any intra-SARS-CoV-2 interactions, the major protein complexes known from the literature were recapitulated: N-N [21]; N-Nsp3 [22]; Nsp7-8-12 [23]; Nsp14-10-16 [24, 25]. We therefore concluded that our SARS-CoV-2 Y2H clone set gave highly specific interactions corroborated by the literature and is thus well suited for Y2H interaction screening.

Next, we cloned a set of human kinases, again in the four Y2H vectors. The kinases in this set were prioritized according to activity measurements in a previous yeast screen [26] which identified active kinases that catalyze phosphorylation-dependent protein interactions. The set comprised a total of 170 protein kinases and 10 non-catalytically active kinase subunits such as cyclins (CCNs) or PRKRA, which were included based on literature (**Suppl. Table 1**). Cyclins are required partner proteins for CDK activation and PRKRA is the protein activator of interferon-induced protein kinase EIF2AK2 (PKR), which was implicated in dsRNA-activated host response to viral infection [27]. The 180 kinase proteins were covered by 200 individual ORFs. Altogether, 368 prey and 377 bait kinase constructs (including the two vector configurations) were used to screen for SARS-CoV-2-human kinase interactions (**Figure 1A**). In this setup, each kinase – SARS-CoV-2 ORF pair was tested in eight configurations (i.e., combinations of different Y2H clone pairs). Testing protein interactions in eight configurations in a stringent Y2H setup strongly increases sensitivity, as most interactions can only be detected in one or few Y2H configurations, while the specificity of the experiment remains unchanged [19, 28, 29]. In total, 56,270 pairs were tested in duplicates utilizing a 384 matrix colony format [30] (**Figure 1A**). As expected, most interactions such as NSP5-PHKG2 or ORF6-FER were identified in only one specific configuration (**Figure 1B**). Noteworthy, we detected the interaction between SARS-CoV-2 ORF9b and MARK2 (microtubule affinity regulating kinase 2) in two configurations (**Figure 1B**), in agreement with the observation that the ORF9b-MARK2 interaction is relatively stable as it has previously been identified in two independent AP-MS studies [3, 4]. In summary, we report a network of 51 interactions involving 14 SARS-CoV-2 ORFs and 29 human kinase proteins (**Figure 1C, Suppl. Table 2**).

**Figure 1:**
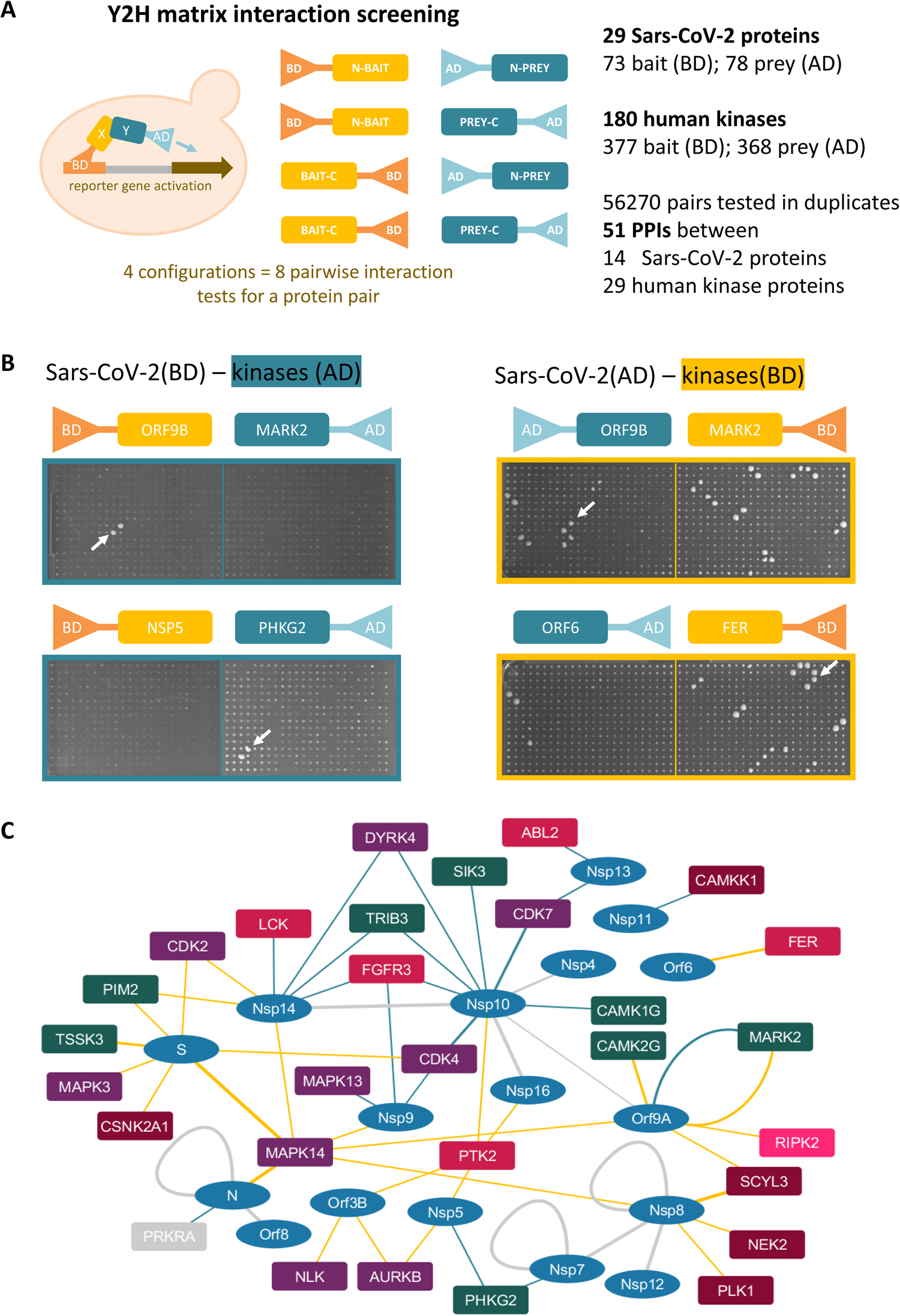
SARS-CoV-2 – human kinase protein interactions. (A) Scheme of Y2H matrix interaction screening. The interaction of 29 SARS-CoV-2 ORFs were screened against 180 human protein kinases in duplicates and 4 configurations against either the binding domain (dark yellow) or the activation domain (petrol blue) matrices. (B) Growth on selective agar: The interaction between SARS-CoV-2 ORF9b and MARK2 (microtubule affinity regulating kinase 2) was detected in two configurations (upper panel). Most interactions such as NSP5-PHKG2 or ORF6-FER were identified in one specific configuration only (lower panel). (C) SARS-CoV-2 – human protein kinase interaction network illustrating 51 interactions between 14 SARS-CoV-2 proteins (ellipsis, blue) and 29 human protein kinases (rectangles colored according to kinase group) determined either by screening the binding domain (dark yellow) or the activation domain (petrol blue) kinase matrices. Interactions between SARS-CoV-2 ORFs shown as grey edges.

The Y2H kinase matrix was a representative subset of the human kinome [31] including members of all major protein kinase groups (**Figure 2A**). Nevertheless, we did not identify any interactions with kinases from the large AGC or STE groups (such as PKA and PKC or MAP2K and PAK, respectively). Conversely, a relatively high number of CAMKs (CAMKs, CAMKLs) and CMGC kinase family members (MAPKs, CDKs) (**Figure 2A**), as well as four non-receptor tyrosine kinases (TK) were found to interact with SARS-CoV-2 proteins.

**Figure 2:**
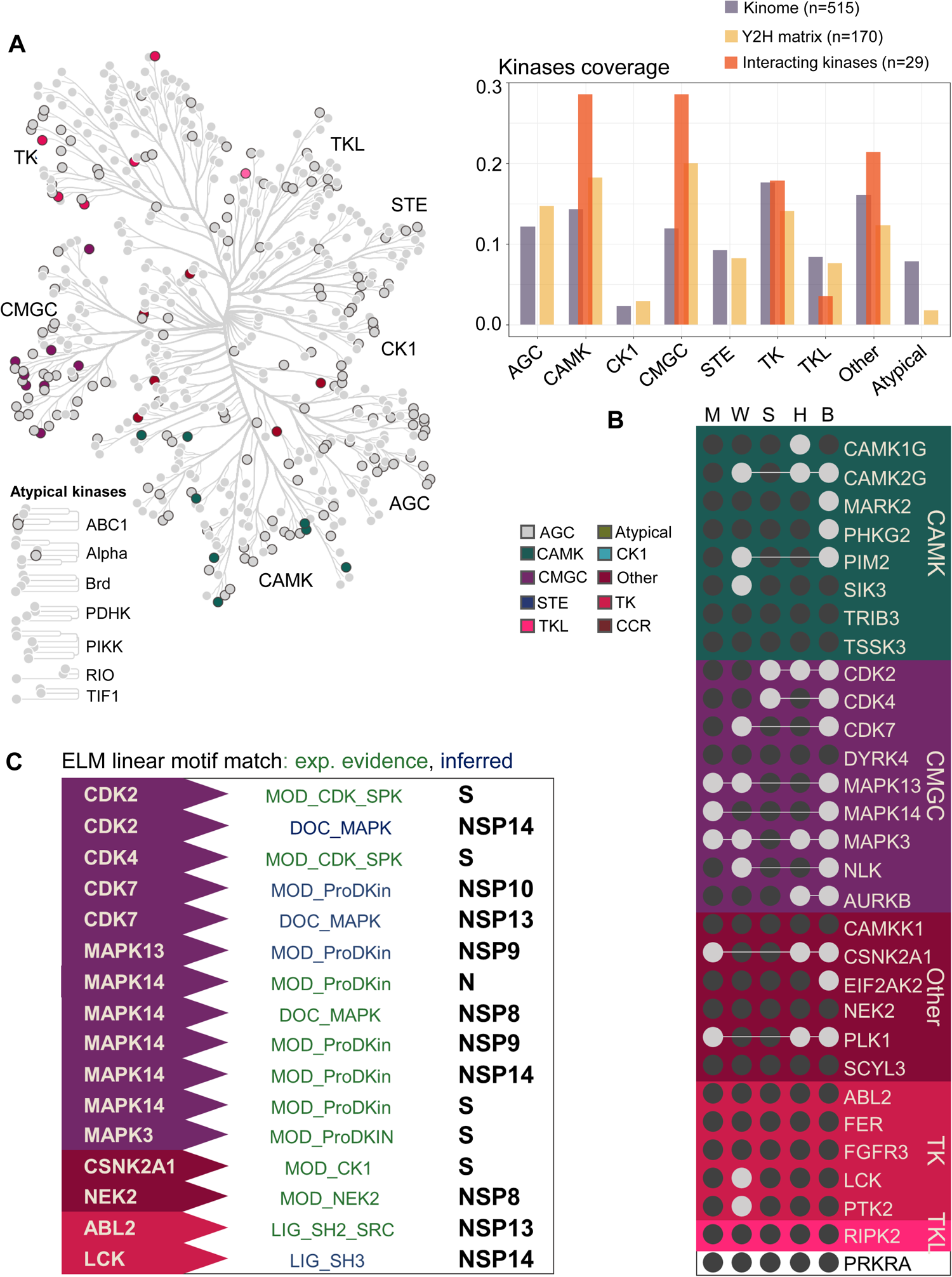
Kinase annotation. (A) Kinome tree with human protein kinases included for the SarsCoV-2 Y2H interaction screen (solid black border). Interacting kinases are colored according to kinase group. Bar chart showing kinase coverage in the Y2H screen: Fraction of human protein kinases across different kinase groups within the whole kinome (gray, n=515), kinases in the Y2H interaction screening (yellow, n=180), and of interacting kinases (orange, n=29). (B) Intersection of interacting kinases and kinases analyzed by previously published surveys: Bouhaddou et al., 2020 (B), Hekman et al., 2020 (H), Stukalov et al., 2021 (S), Mari et al., 2022 (M), Wang et al., 2022 (W), colored by kinase group. (C) Inferred (blue) Eukaryotic Linear Motifs (ELM) or with exp. evidence (green) predicted to contribute to the interaction between human kinases and SARS-CoV-2 proteins.

Previously, the changes in protein phosphorylation upon SARS-CoV-2 infection of cultured cells have been determined by phospho-proteomics [4, 9, 10, 12, 13]. Kinase-substrate sets, assembled from the literature, were then used to query those datasets for kinase activity patterns through enrichment analyses (kinase-substrate enrichment analysis, KSEA; [32]). When we directly compared kinase activity changes inferred from these phospho-proteomics studies, we found that 60% (18/28) of the kinases that interacted with SARS-CoV-2 proteins in our study showed a change in activity profile in one or more studies (**Figure 2B**). Moreover, one of the two kinase families most enriched in our screen, the CMGC family which encompasses MAPK and CDK kinases, has also been previously proposed to account for a substantial proportion of the phospho-proteome response upon SARS-CoV-2 infection both in human cells and in mice [9, 12].

In contrast to AP-MS data, the vast majority of Y2H interactions represent direct, binary interaction information [16]. Therefore, we performed a linear motif analysis to identify putative interaction interfaces and phosphorylation sites in SARS-CoV-2 proteins that interact with human kinases (**Figure 2C**). We used the *mimic*INT computational workflow [33] to computationally infer and score interaction interfaces between SARS-CoV-2 and human proteins mediated by motif-domain interaction templates from the ELM database [34]. The *mimic*INT scoring system consists of two metrics. First, a similarity score between the domain of the interacting protein and a domain that had been previously known to bind to a given ELM motif was calculated. Second, the probability of a given motif to occur by chance in a viral sequence is statistically assessed by randomizing its disordered regions 100,000 times [35]. To include an additional layer of specificity in the inference, we used the Automatic Kinase-specific Interaction Detection (AKID) method [36] that exploits both the sequence of the putative substrate, as well as the kinase specificity determinants to infer kinase-substrate interactions. Combining the MimicINT motif approach with a stringent AKID score (threshold > 0.7), 16 interactions were modelled successfully (**Figure 2C, Suppl. Table 3**). For example, we identified a S-CSNK2A1 phosphorylation motif (MOD_CK1_1) driven interaction supported by a high kinase domain similarity score (1.2), statistical significance in motif occurrence (p = 0.0055) and a literature reported phospho-site candidate (S816). In general, phosphorylation motifs such as those from proline-directed kinases (CDKs and MAPKs; MOD_ProDKin_1) dominated the results, often providing more than one motif match per interaction. A MAPK docking motif (DOC_MAPK_gen_1) interaction supported the NSP8-MAPK14 interaction. The interactions NSP13-ABL2 and NSP14-LCK were predicted to be mediated by SH2 (LIG_SH2_SRC) and SH3 (LIG_SH3_3) domain linear motif recognition, respectively. This *in silico* identification of putative motif mediated interaction interfaces and phosphorylation sites in SARS-CoV-2 proteins adds further evidence to support the novel interactions identified in our screen.

Next, we aimed to leverage the extensive number of RNA-seq datasets reporting on expression changes upon SARS-CoV-2 infection. Recently, the data were normalized and aggregated in the COVID19db platform [37]. Changes in a kinase activity are reflected in the transcriptional activity of the cell. To infer the alterations in an activity of a kinase, the Kinase Enrichment Analysis (KEA) uses prior knowledge about expression changes of sets of genes coregulated with the kinase. The Ma’ayan Lab updated their kinase enrichment analysis workflow [38] and computed two distinct Kinase co-expression libraries, each covering co-expression signatures for most human kinases. A signature is built form the top 300 genes whose expression is most significantly correlated with the expression of a human kinase across one of two large gene expression data sets, namely GTEx or ARCHS4 [39, 40].

We hypothesized that a direct SARS-CoV-2 protein interaction with a human kinase may alter the kinase activity. To test whether we can assess these changes in kinase activity based on transcriptomic data, we looked for kinase specific gene set enrichment [41] in transcriptome data from kinase-loss of function (LoF) or kinase knockout cell lines (A549, A375, H1AE, HEPG2, HT29, MCF7, and PC3) of the core CMap expression compendium [42]. Indeed, we found statistically significant positive or negative normalized enrichment scores (NES) with both the GTEx or ARCHS4 kinase activity gene set in more than half of the respective kinase-LoF cell lines (**Figure 3A**). For example, AURKB, CDK4, CDK7, FER, and MAPK13 showed significant negative NES, while CDK2, CAMK2G, NEK2, and RIPK2 exhibited positive NES, for both GTEx- and ARCHS4-derived query kinase signatures across multiple kinase-LoF cell lines.

**Figure 3:**
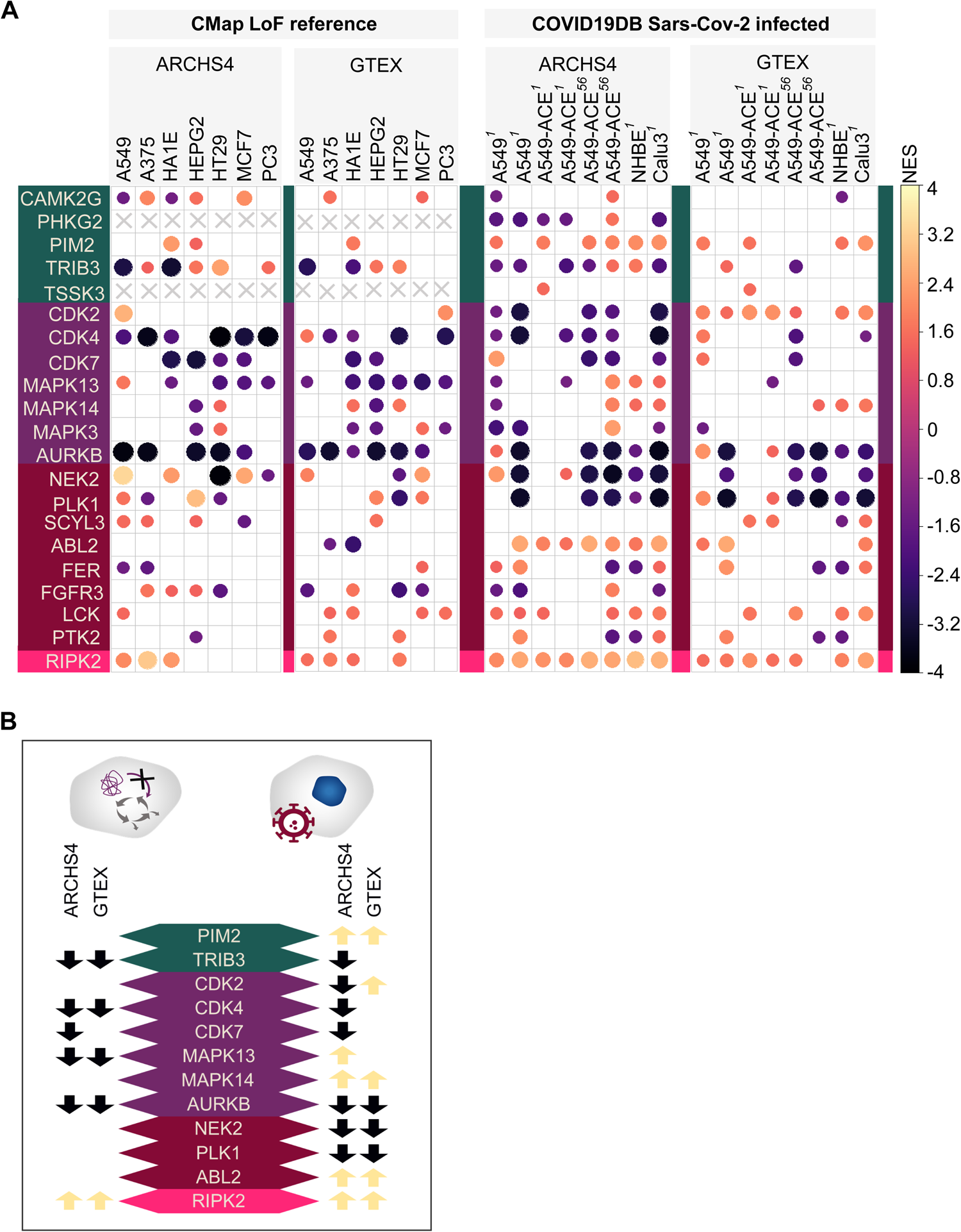
Transcriptional changes associated with human kinases. (A) Normalized enrichment scores (NES) of significantly (adjusted p-value ≤ 0.05) positively or negatively enriched kinases tested either in the ARSCHS4 or GTEX co-expressed gene sets for either the reference CMap loss of function (LoF) expression sets of each human protein kinase (left panel) or for publicly available RNA-Seq data of infected SARS-CoV-2 cell lines downloaded from Covid19db (right panel). Numbers in superscript represent the corresponding COVID10db id. (B) Summary of enrichment analysis. Black arrows encode for negative enrichment of the kinase in the respective condition throughout cell lines, and bright arrows for positive enrichment. Congruent enrichment between LoF and SARS-CoV-2 infected conditions indicate to inhibition of the particular kinase.

We next took RNA-seq transcriptome data from SARS-CoV-2 infection models, prioritizing the A549 lung cancer cell lines, as they were extensively used in experiments monitoring changes in gene expression upon infection [37, 43]. We again calculated enrichment scores for the two kinase co-expression libraries. In analogy to what we had observed with kinase-LoF cell lines, significant positive or negative enrichments indicative of altered kinase activity, were found in several SARS-CoV-2 A549 infection datasets. ABL2, LCK, MAPK14, PIM2, and RIPK2 showed positive NES across multiple RNA-seq datasets for both GTEx- and ARCHS4-derrived kinase signatures, whereas CDK4, CDK7, CSNK2A1, FER, MAPK3, NEK2, PHKG2, PLK1, and TRIB3 kinase sets showed negative NES (**Figure 3A**). While those results pointed to a change in activity during a SARS-CoV-2 infection for several kinases, it is not straightforward to conclude whether these findings indicated an increase or decrease in the activity of a given kinase. However, since we had previously assessed the kinase-LoF profiles from CMap cell lines, we could identify a set of kinases with the same sign of NES scores in both the kinase-LoF and the SARs-Cov-2 RNA-seq data, whether negative or positive, and another group with opposite signs in the two analyses (**Figure 3B**). We therefore postulate that kinases with concordant NES signs (that is the same NES in the infection context as in the kinase-LoF cell line) have a decreased activity upon SARS-Cov-2 infection. Conversely, the opposite NES signs may then point towards increased kinases activity (**Figure 3B**). The activity alterations of twelve kinases with a clear NES pattern from the two analyses were summarized in **Figure 3B**, depicting SARS-CoV-2-dependent kinase inhibition of AURKB, CDK4, CDK7, and RIPK2, and activation of MAPK13.

In addition to transcriptomics, several phospho-proteomics data sets were recorded from SARS-CoV-2 infected cell models. Therefore we also directly probed the phosphorylation response for kinase activity changes [13, 38, 44]. While quantitative changes of phosphorylation site abundance offer a more direct measure of kinase activity than RNA expression profiling, the vast majority of phospho-sites are not assigned to any protein kinase, leading to relatively small substrate sets sizes. Consequently, the analysis is often limited due to the restricted overlap between query protein sets and observed phospho-sites peptides.

We employed an established kinase substrate set for kinase activity profiling through enrichment analyses [44] against the ranked phosphorylation data sets of a SARS-CoV-2 infection. Of the 29 kinases in our interaction data set, eleven had 20 or more annotated substrates (**Figure 4A**). We found that CDK7, CDK2, CDK4, MAPK14, MAPK13, MAPK3, AURKB, and PLK1 substrate patterns showed a significant phospho-protein change upon SARS-CoV-2 infection (**Figure 4B**).

**Figure 4:**
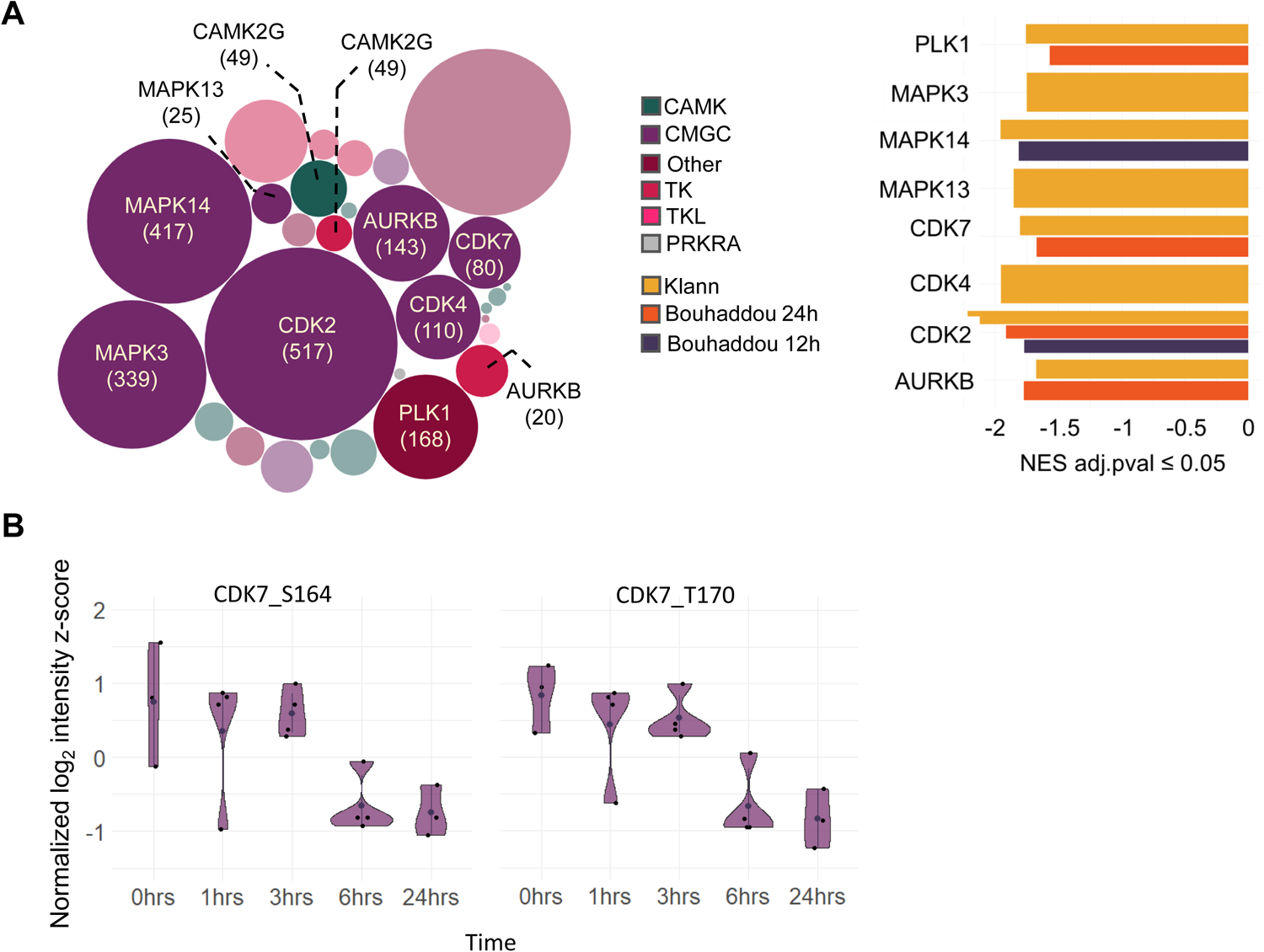
Phosphorylation changes associated with human kinases. (A) Overview of used substrate sets for each kinase (number of substrates per kinase in brackets) used for enrichment analysis, colored by kinase group. Kinases excluded from enrichment analysis are not annotated and shown as semi-transparent bubbles. (B) Normalized enrichment scores (NES) of significantly enriched (adjusted p-val <=0.05) kinases in either the phosphorylation data of Klann et al. in yellow [11], or Bouhaddou et al. for 12h (purple) and 24h (orange) [9]. (C) Phosphorylation changes for CDK7_S164, and CDK7_T170 reported by Heckmann et al. [10] for four different timepoints with a strong decrease at 6h.

Kinase activity in cells is also reflected by the phosphorylation status of the kinase itself, such as the phosphorylation of the conserved kinase activation loop [45, 46]. Several of the kinases that directly interact with SARS-CoV-2 proteins in our network, including ABL2, CAMK2G, CDK2, CDK7, MAPK3, MAPK13, MAPK14, PLK1, PTK2, RIPK2, SIK3, and PRKRA, are themselves differentially phosphorylated upon SARS-CoV-2 infection [4, 9–13]. The most consistent and prominent effect was a decrease of the CDK7 activation loop phosphorylation (at S164 and T170) upon SARS-CoV-2 infection in three data sets [4, 9, 10]. Notably, Hekmann et al. (2020) measured a strong decrease of T170 and S164 phosphorylation 6h post infection (**Figure 4C**). CDK7 functions catalytically in a cycle with CDK2 through phosphorylation of the T-loop [47] and a T170A substitution results in a total loss of CDK7 activity [48]. This result parallels our transcriptome data analyses, as both analyses indicated decreased CDK7 activity upon SARS-CoV-2 infection.

To demonstrate a direct modulation of a kinase activity through interaction with a SARS-CoV-2 protein, we focused on a novel, previously unreported interaction between human tyrosine kinase FER and SARS-CoV-2 ORF6 (**Figure 1C**). ORF6 mutation and expression status is critical for the pathological impact of the various SARS-CoV-2 strains, making ORF6 and its downstream effects prime targets of therapies aiming at reducing the disease severity [14, 49].

Based on our previous work where we assayed human kinase activities in yeast [26, 50, 51], we used a yeast system to characterize this interaction in more detail. Human tyrosine kinases activity, when exogenously expressed in yeast, can be assayed through measuring growth effects or the phosphorylation of yeast proteins through western blotting [52]. Importantly, *Saccharomyces cerevisiae* itself hardly has any intrinsic phospho-tyrosine kinase activity [51], so that all detected phospho-tyrosines can be unambiguously attributed to the exogenously expressed human kinase. To investigate the effect of ORF6 on the FER kinase activity, we expressed full length human FER in yeast (Saccharomyces cerevisiae L40c [30]) in combination with ORF6, other SARS-CoV-2 control proteins, or an empty control (**Figure 5**). Upon expression of FER kinase phospho-tyrosine activity was detected through immuno-blotting with the 4G10 pan phospho-tyrosine antibody. In the yeast strains that were transformed with an ORF6 construct, FER activity was strongly reduced, suggesting that co-expression and interaction with ORF6 inhibits FER kinase activity in living yeast. Expression of other, non-interacting SARS-CoV-2 proteins had no effect on the phosphorylation levels of yeast proteins. Yeast growth, and total protein amount were not different between ORF6-FER and control-FER strains.

**Figure 5:**
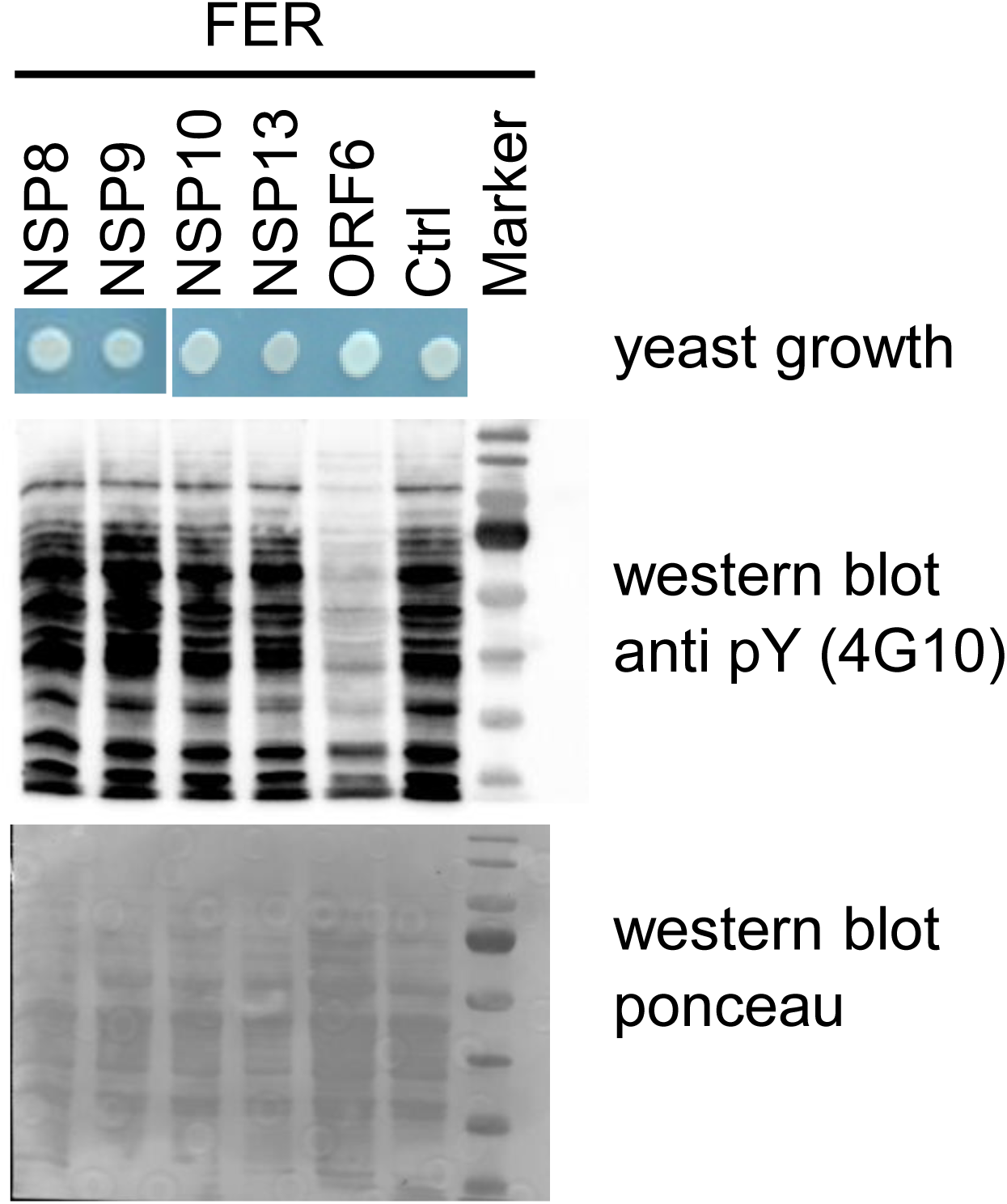
A SARS-CoV-2 – host interaction affects FER tyrosine kinase activity. Top: Yeast growth on selective agar, expression of FER together with SARS-CoV-2 proteins in yeast does not impair growth. Middle: Anti-pY (4G10) western blot immunostaining of phosphorylated yeast proteins in full cell lysates. A yeast strain expressing ORF6 together with human FER shows strongly reduced anti-pY AB staining of yeast proteins. Bottom: Loading control of the full yeast soluble protein lysate through ponceau staining of the membrane.

ORF6 is among the most pathogenic proteins encoded by the SARS-CoV-2 genome, exhibiting highest cytotoxicity when overexpressed in HEK293T [49]. It modulates the relative expression levels of the other SARS-CoV-2 proteins, both structural and accessory, via posttranscriptional mechanisms. In addition, ORF6 harbors a predicted membrane-embedded alpha-helix motif on its N-terminus and has been detected at the peripheral membrane [53], in the endoplasmic reticulum and at the membranes of a subset of the intracellular vesicles, including autophagosome and lysosomes [49], as well as at lipid droplets [54].

ORF6 has been found to inhibit type I interferon (IFN) production and signaling [55, 56], at least in part by blocking the nuclear import of the transcription factors STAT1, STAT2, and IRF3 either through direct interaction or through binding to the nuclear pore complex protein Nup98 [55, 57]. In strains with high levels of ORF6 interferon response genes are suppressed. It also inhibits the nuclear export of mature mRNAs, leading to an impairment of the host antiviral responses [53, 58].

ORF6 is prominently involved in suppressing the innate immune response to the virus. Suppression of innate immune response caused by the different SARS-CoV-2 variants correlated with ORF6 expression levels of the virus variants. Specifically, comparison of the SARS-CoV-2 variants of concern (such as alpha, delta, or omicron) revealed that the balance of adaptive and innate immune escape during the course of strain evolution is linked to ORF6 mutations and protein expression level [14].

Intriguingly, FER was demonstrated to be involved in innate immune response as well. FER phosphorylates one of the key transcription factors in innate immune response, STAT3, at Y705, which leads to its dimerization and nuclear translocation and activation of its target genes [59, 60]. Through STAT activation, enhancing innate immunity, FER overexpression improved survival of a C57/BL6 mice pneumonia models accelerating bacterial clearance in the lung [61]. There are interesting parallels with the original SARS virus, where decreased phosphorylation of STAT3 on Y705 was observed, leading to increased viral replication [62].

Finally, a recent mechanistic study demonstrated that human tyrosine kinase FER regulates endosomal recycling. FER depletion or inhibition of its kinase activity leads to an impairment of Rab4-recycling endosome formation in inducible FER knockdown MDA-MB-231 cells [63]. The endosomal entry pathway, as one of two distinct mechanisms for SARS-CoV-2 cell entry [64], involves S protein binding and internalization via the cathepsin-dependent endosomal pathway [65], where FER may be involved. These observations suggest possibilities for a cross-talk between the two interacting proteins, ORF6 and FER, guiding further research into their role during SARS-CoV-2 infection.

## Conclusion

In this study we define SARS-CoV-2 protein interactions with human kinases. Because of its transient nature, protein interactions with human kinases are often underrepresented in large scale data sets, such as those generated for SARS-CoV-2 viral host interactions previously. Exhaustive Y2H screening of all SARS-CoV-2 proteins against a set of 180 human kinase proteins revealed 51 potential interactions between 14 viral proteins and 29 kinase proteins (**Figure 1**). A previously well-characterized physical interaction between ORF9A and MARK2 [14], as well as a less characterized, direct interaction between protein N and PRKRA [8] were recapitulated in our screen.

The widespread phospho-response upon SARS-CoV-2 infection, coupled with the availability of a large range of kinase inhibitors, has sparked investigations into the potential utilization of these compounds as antiviral, anti-inflammatory, cytokine-suppressive, or antifibrotic treatments in SASRS-CoV-2 infections [66, 67]. In our systematic interaction study, we specify direct interactions between SARS-CoV-2 proteins and 29 human kinases, with a relatively high number of interactions for CAMK, CMGC and non-RTK kinase family members. This is in agreement with results based on phospho-proteomic experiments monitoring the course of SARS-CoV-2 infection in cells [9, 10], where an involvement of p38 MAPKs (MAPK14), Erk (MAPK3), CDKs (such as CDK2), CK2, and SRPKs, was inferred indirectly through activity profiling (**Figure 2**). p38 MAPK and Erk were found to be activated during SARS-CoV-1 infection as well [62]. In addition, CDKs play key roles in controlling viral replication [68], not only through their cellular functions in the cell cycle but also by acting on SARS-CoV-2 proteins directly [21]. Our data provide additional support for a SARS-CoV-2 dependent inhibition of the transcriptional kinase CDK7. We report novel evidence that ORF6, through physical interaction, can inhibit tyrosine kinase FER activity (**Figure 5**). This is corroborated by literature reports that show that both ORF6 and FER impact STATs, the key transcription factors in innate immune response, in opposite directions. Therefore, a valid hypothesis for further investigation is that the ORF6 repressive effect on STAT-mediated innate immune response may in part be related to its ability to inhibit FER kinase activity.

## Materials and Methods

### Clone construction

Two independent sets of clones were used, to complement each other for missing sequences. And additionally, to take advantage of different linker sequences efficacy to increase sensitivity (at the same specificity) [29].

First, we obtained a copy of the mammalian expression clones for PCR subcloning into our gateway compatible cloning system, into the entry vector pDONR221 (Thermofisher) from the Krogan lab at UCSF, USA, [3] using the following primers (C19_rev: ACCACTTTGTACAAGAAAGCTGGGTCtcccccgccgccttcgag and C19_fwd: ACAAGTTTGTACAAAAAAGCAGGCTTCggtgaattcgccgccacc). The set is lacking the large NSP3 and NSP16 protein sequences. Second, the set of SARS-CoV-2 ORF clones were obtained as a courtesy of the labs of F.P. Roth (Toronto) and Y. Jacob (Paris) [20] This set includes all ORFs, except the very small NSP11, ready in Gateway cloning compatible entry vectors pDONR223 [69] or pDONR207 (Invitrogen, USA) respectively. The GATEWAY re-combinatorial system (Invitrogen) is used to shuttle the two sets of SARS-CoV-2 ORFs into the pBTM116-D9 [N-lexABD] and pBTMcC24-DM [C-lexABD] bait and pACT4-DM [N-Gal4AD] and pCBDU-JW [C-Gal4AD] prey vectors, respectively [30].

### Y2H protein-protein interaction screening

The Y2H matrix screen was performed as described previously [30]. L40c MAT**a** and Mat**α** yeast strains were transformed with the bait and prey plasmids, respectively, using the lithium acetate method in 96-well format and finally stamped on selective NB-agar using a gridding robot (Kbiosystems Limited, UK) and incubated at 30 °C for 3-4 days.

The Y2H mating approach was performed in 384 microtiter-plate format including two biological replicates (independently transformed yeast colonies) per Y2H bait-prey vector pair. Colony growth on SD5 agar plates lacking leucine, tryptophan, adenine, uracil and histidine after 5 days at 30 °C (see **Figure 1**) was visually assessed for protein-protein interaction detection. Only Y2H pairs that grew in both replicates were considered in the analyses.

### Assaying FER kinase activity in yeast

The yeast strains L40c Mat**α** was co-transformed with SARS-CoV-2 ORF6 or control proteins (pRS425-GPD) and the FER kinase (P16591 in pASZ-DM) and grown in 2 ml selective media including 100 µM CuSO_4_ at 30 °C oN. Cells were lysed for 10 min on ice in 250µl 1M NaOH/ 7.5% β-mercaptoethanol. Then 250 µl 50% TCA was added and incubated for another 10 min on ice. Cells were pelleted through centrifugation at 4 °C at 15000 rpm. The pellet was washed twice with MQ water and resuspended in SDS-gel loading buffer. Samples were separated using 10% SDS page and blotted on a nitrocellulose membrane using the Trans-Blot Turbo Transfer System (Bio-Rad Laboratories, Inc., Austria). The membrane was probed with the anti pY-4G10 (mouse, Millipore, 05-321), the secondary mAB anti-mouse (sheep, GE Healthcare, LNA931V) and developed using Clarity Western ECL Substrate (Bio-Rad Laboratories, 1705061) and the UVP ChemStudio (Analytik Jena GmbH, Germany) to record the chemiluminescence signal.

### Kinase-substrate interaction motif analysis

The *mimic*INT computational interaction inference workflow[33] is based on sequence and structural data analysis and allows to computationally identify interaction interfaces mediated by short linear motifs (pre-defined by ELM database [34]). A similarity score between the domains of the human interacting protein to a domain that was known to bind a given ELM motif is calculated [35]. The putative motif in the viral protein sequence is empirically assessed for statistical significance against the distribution of the number of ELM motif occurrences. Briefly, we randomized 100,000 times the disorder regions in the viral protein sequences. Disorder propensity is predicted using the IUPred algorithm [70] (randomizations were performed on disordered regions using the 0.2 disorder score threshold with long disorder option), and disorder regions are shuffled with two different strategies [71] (i) shuffling the disorder regions of each sequence individually, and (ii) shuffling the disorder regions across all sequences. This allowed the generation of two disorder background sets for the determination of statistical significance of the motif matches between SARS-CoV-2 linear sequences and the human kinases. 22 interactions are supported by at least one predicted linear motif interaction site.

Automatic Kinase-specific Interaction Detection (AKID) method [36] exploits the sequence of the putative substrate, as well as the kinase specificity determinants to infer kinase-substrate interactions as phospho-sites. We combined the two approaches and taking sites with a stringent AKID score > 0.7 as additional specificity criteria for interaction prediction (**Suppl. Table 3**).

### Data collection and processing

Reference expression profiles for each kinase were downloaded as loss of function (LoF) consensus signatures (moderated zscore: MODZ) for all available cell lines from Connectivity Map (CMap, LINCS 2020). To ensure data homogeneity only datasets with a perturbation time of 96 h and perturbation dose of either 1 or 1.5 were considered for the seven most frequent core cell lines (A549, A375, HA1E, HEPG2, HT29, MCF7, PC3). This resulted in a final set of 23 kinase-LoF profiles for enrichment analysis, see **Suppl. Table 4**.

SARS-CoV-2 expression signatures were obtained as standardized and normalized transcriptomic data from the COVID19db database platform [37]. 95 data sets and its corresponding metadata were downloaded and manually filtered. Non-SARS-CoV-2, drug treated or data sets missing healthy controls have been excluded from further processing and analysis. Transcriptomic data sets were manually assigned to single experiments and SARS-CoV-2 infected and healthy controls according to the provided Covid19db metadata for each entry. A custom R script implementing the Bioconductor DESeq2 package [72] was used to assess differential changes between SARS-CoV-2 and healthy controls across different cell lines, cell types, and tissues according to the latest documentation at https://bioconductor.org/packages/release/bioc/vignettes/DESeq2/inst/doc/DESeq2.html.

Two co-expression sets, ARCHS4 and GTEX, comprising top 300 co-regulated genes, for each kinase, were acquired from the Ma’ayan lab to perform enrichment analysis using the fgsea R package [73] according to its latest documentation. First, enrichment analysis was carried out for the reference CMap kinase-LoF consensus signatures, and second for the previously calculated expression profiles from the COVID19db data. In each case, descending sorted expression sets (log fold change (LFC): COVID19db, moderate zscore: CMap) were used as rank sets in combination with either the ARCHS4 or the GTEX enrichment set for the corresponding kinase to assess the normalized enrichment scores (NES) and its significance. Whenever gene names/symbols were used, these were converted into the corresponding EntrezIds according to the HUGO Gene Nomenclature Committee (HGNC) complete set (version April 2021). Entries with no matching ID have been discarded from further analysis.

Enrichment analysis was repeated similarly using phospho-proteomic data previously published by Stukalov et al. 2020 [4], Klann et a. 2020 [11], Bouhaddouh et al. 2020 [9], and Hekman et al.2020 [10] as rank sets and an established substrate set from Dogourd et al. 2021 [44].

### Data visualization

Tailored R scripts implementing ggplot2 [74], corrplot (Wei T, Simko V (2021). *R package ‘corrplot’: Visualization of a Correlation Matrix*. (Version 0.92), https://github.com/taiyun/corrplot), and bubbles were used for data visualization if not stated otherwise. Interaction networks have been created using Cytoscape [75]. Coral [76] was used to visualize target kinase distribution on the kinome tree.

## Author Information

These authors contributed equally: Bettina Halwachs, Christina S. Moesslacher, Johanna M. Kohlmayr;

conceptualization: US;

methodology: US, BH, CSM, JMK;

software: BH, AZ, SAC;

clone-construction: CSM, JMK, SF;

yeast-to-hybrid screening: CSM, JMK, SF;

formal analysis: BH, US, AZ;

investigation: BH, CSM, JMK, SF, SM, SAC, AZ, US;

resources: US, AZ;

data curation: US, BH, JMK, CSM;

writing: US, BH, NK;

visualization: BH, CSM, JMK, US;

supervision, administration, and funding: US;

All authors discussed, read, and commented on the manuscript.

## Acknowledgments

We thank N.J. Krogan, F.P. Roth, and Y. Jacob for providing plasmids with SARS-CoV-2 ORFs. Centre de Calcul Intensif d’Aix-Marseille is acknowledged for granting access to its high performance computing resources.

## Supplementary material

**Supplemental Figure 1:**
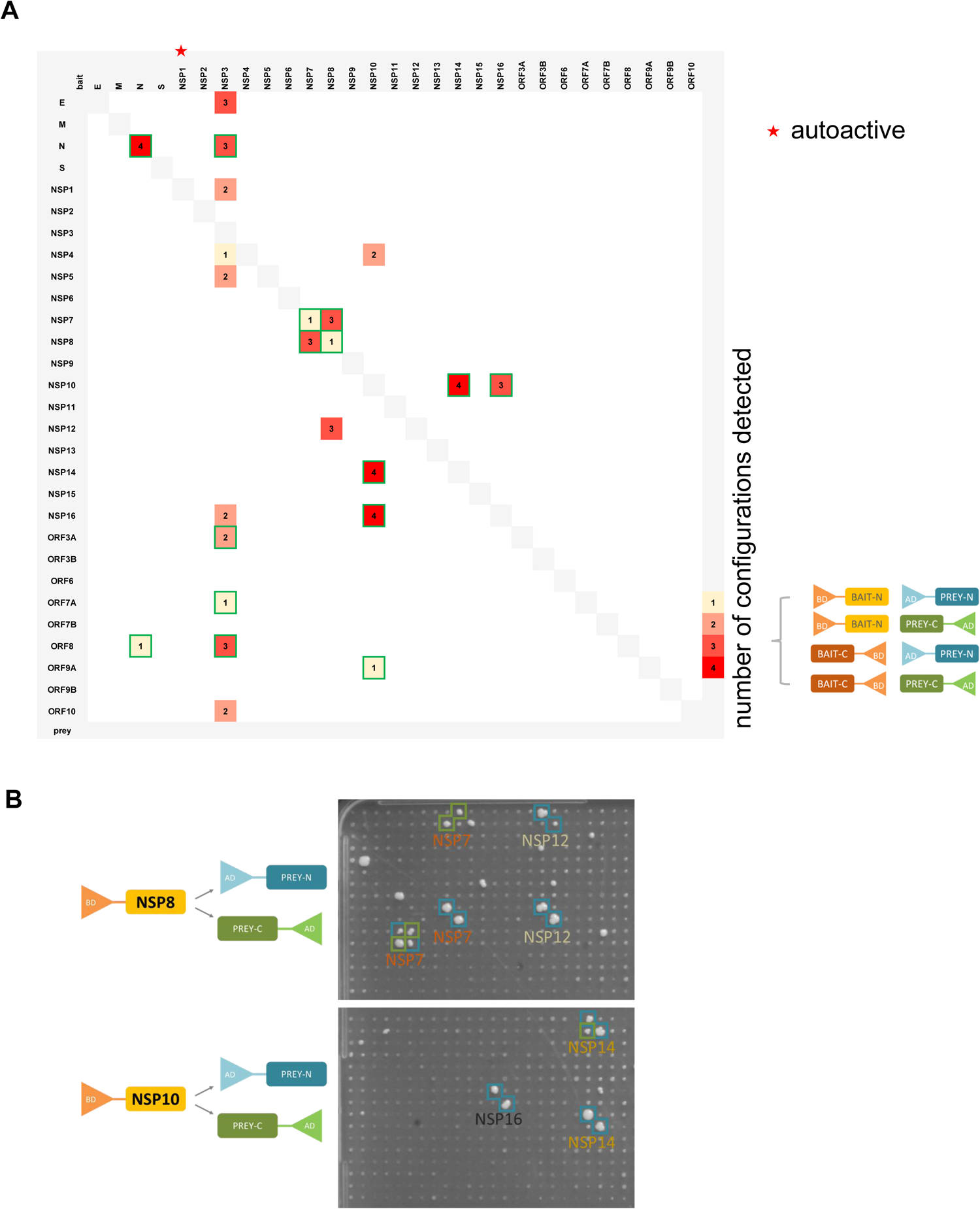
SARS-CoV-2 – SARS-CoV-2 viral protein interactions. (A) Matrix depicting protein-protein interaction detected through Y2H analysis of SARS-CoV-2 ORFs. (B) Growth on selective agar: examples of Y2H results for the NSP8 and NSP10 proteins when used as N-terminal lexADB-fusion proteins.

## References

1. Wu F, Zhao S, Yu B, Chen Y-M, Wang W, Song Z-G, et al. A new coronavirus associated with human respiratory disease in China. Nature. 2020;579:265–9. doi:10.1038/s41586-020-2008-3.

2. Varona JF, Landete P, Lopez-Martin JA, Estrada V, Paredes R, Guisado-Vasco P, et al. Preclinical and randomized phase I studies of plitidepsin in adults hospitalized with COVID-19. Life Sci Alliance 2022. doi:10.26508/lsa.202101200.

3. Gordon DE, Jang GM, Bouhaddou M, Xu J, Obernier K, White KM, et al. A SARS-CoV-2 protein interaction map reveals targets for drug repurposing. Nature. 2020;583:459–68. doi:10.1038/s41586-020-2286-9.

4. Stukalov A, Girault V, Grass V, Karayel O, Bergant V, Urban C, et al. Multilevel proteomics reveals host perturbations by SARS-CoV-2 and SARS-CoV. Nature. 2021;594:246–52. doi:10.1038/s41586-021-03493-4.

5. Liu X, Huuskonen S, Laitinen T, Redchuk T, Bogacheva M, Salokas K, et al. SARS-CoV-2-host proteome interactions for antiviral drug discovery. Mol Syst Biol. 2021;17:e10396. doi:10.15252/msb.202110396.

6. Li J, Guo M, Tian X, Wang X, Yang X, Wu P, et al. Virus-Host Interactome and Proteomic Survey Reveal Potential Virulence Factors Influencing SARS-CoV-2 Pathogenesis. Med (N Y). 2021;2:99–112.e7. doi:10.1016/j.medj.2020.07.002.

7. Zhou Y, Liu Y, Gupta S, Paramo MI, Hou Y, Mao C, et al. A comprehensive SARS-CoV-2-human protein-protein interactome reveals COVID-19 pathobiology and potential host therapeutic targets. Nat Biotechnol. 2023;41:128–39. doi:10.1038/s41587-022-01474-0.

8. Kim D-K, Weller B, Lin C-W, Sheykhkarimli D, Knapp JJ, Dugied G, et al. A proteome-scale map of the SARS-CoV-2-human contactome. Nat Biotechnol. 2023;41:140–9. doi:10.1038/s41587-022-01475-z.

9. Bouhaddou M, Memon D, Meyer B, White KM, Rezelj VV, Marrero MC, et al. The Global Phosphorylation Landscape of SARS-CoV-2 Infection. Cell 2020. doi:10.1016/j.cell.2020.06.034.

10. Hekman RM, Hume AJ, Goel RK, Abo KM, Huang J, Blum BC, et al. Actionable Cytopathogenic Host Responses of Human Alveolar Type 2 Cells to SARS-CoV-2. Mol Cell. 2020;80:1104–1122.e9. doi:10.1016/j.molcel.2020.11.028.

11. Klann K, Bojkova D, Tascher G, Ciesek S, Münch C, Cinatl J. Growth Factor Receptor Signaling Inhibition Prevents SARS-CoV-2 Replication. Mol Cell. 2020;80:164–174.e4. doi:10.1016/j.molcel.2020.08.006.

12. Wang ZN, Yang XS, Sun J, Zhao JC, Zhong NS, Tang XX. Multi-omics evaluation of SARS-CoV-2 infected mouse lungs reveals dynamics of host responses. iScience. 2022;25:103967. doi:10.1016/j.isci.2022.103967.

13. Mari T, Mösbauer K, Wyler E, Landthaler M, Drosten C, Selbach M. In Vitro Kinase-to-Phosphosite Database (iKiP-DB) Predicts Kinase Activity in Phosphoproteomic Datasets. J Proteome Res 2022. doi:10.1021/acs.jproteome.2c00198.

14. Bouhaddou M, Reuschl A-K, Polacco BJ, Thorne LG, Ummadi MR, Ye C, et al. SARS-CoV-2 variants evolve convergent strategies to remodel the host response. Cell 2023. doi:10.1016/j.cell.2023.08.026.

15. Luck K, Kim D-K, Lambourne L, Spirohn K, Begg BE, Bian W, et al. A reference map of the human binary protein interactome. Nature. 2020;580:402–8. doi:10.1038/s41586-020-2188-x.

16. Kunowska N, Stelzl U. Decoding the cellular effects of genetic variation through interaction proteomics. Curr Opin Chem Biol. 2022;66:102100. doi:10.1016/j.cbpa.2021.102100.

17. Woodsmith J, Stelzl U. Studying post-translational modifications with protein interaction networks. Curr Opin Struct Biol. 2014;24:34–44. doi:10.1016/j.sbi.2013.11.009.

18. Vinayagam A, Stelzl U, Wanker EE. Repeated two-hybrid screening detects transient protein–protein interactions. Theor Chem Acc. 2010;125:613–9. doi:10.1007/s00214-009-0651-8.

19. Venkatesan K, Rual J-F, Vazquez A, Stelzl U, Lemmens I, Hirozane-Kishikawa T, et al. An empirical framework for binary interactome mapping. Nat Methods. 2009;6:83–90. doi:10.1038/nmeth.1280.

20. Kim D-K, Knapp JJ, Kuang D, Chawla A, Cassonnet P, Lee H, et al. A Comprehensive, Flexible Collection of SARS-CoV-2 Coding Regions. G3 (Bethesda). 2020;10:3399–402. doi:10.1534/g3.120.401554.

21. Carlson CR, Asfaha JB, Ghent CM, Howard CJ, Hartooni N, Safari M, et al. Phosphoregulation of Phase Separation by the SARS-CoV-2 N Protein Suggests a Biophysical Basis for its Dual Functions. Mol Cell. 2020;80:1092–1103.e4. doi:10.1016/j.molcel.2020.11.025.

22. Hurst KR, Koetzner CA, Masters PS. Characterization of a critical interaction between the coronavirus nucleocapsid protein and nonstructural protein 3 of the viral replicase-transcriptase complex. J Virol. 2013;87:9159–72. doi:10.1128/JVI.01275-13.

23. Hillen HS, Kokic G, Farnung L, Dienemann C, Tegunov D, Cramer P. Structure of replicating SARS-CoV-2 polymerase. Nature. 2020;584:154–6. doi:10.1038/s41586-020-2368-8.

24. Krafcikova P, Silhan J, Nencka R, Boura E. Structural analysis of the SARS-CoV-2 methyltransferase complex involved in RNA cap creation bound to sinefungin. Nat Commun. 2020;11:3717. doi:10.1038/s41467-020-17495-9.

25. Ma Y, Wu L, Shaw N, Gao Y, Wang J, Sun Y, et al. Structural basis and functional analysis of the SARS coronavirus nsp14-nsp10 complex. Proc Natl Acad Sci U S A. 2015;112:9436–41. doi:10.1073/pnas.1508686112.

26. Jehle S, Kunowska N, Benlasfer N, Woodsmith J, Weber G, Wahl MC, Stelzl U. A human kinase yeast array for the identification of kinases modulating phosphorylation-dependent protein-protein interactions. Mol Syst Biol. 2022;18:e10820. doi:10.15252/msb.202110820.

27. Balachandran S, Roberts PC, Brown LE, Truong H, Pattnaik AK, Archer DR, Barber GN. Essential Role for the dsRNA-Dependent Protein Kinase PKR in Innate Immunity to Viral Infection. Immunity. 2000;13:129–41. doi:10.1016/S1074-7613(00)00014-5.

28. Chen Y-C, Rajagopala SV, Stellberger T, Uetz P. Exhaustive benchmarking of the yeast two-hybrid system. Nat Methods. 2010;7:667–8; author reply 668. doi:10.1038/nmeth0910-667.

29. Choi SG, Olivet J, Cassonnet P, Vidalain P-O, Luck K, Lambourne L, et al. Maximizing binary interactome mapping with a minimal number of assays. Nat Commun. 2019;10:3907. doi:10.1038/s41467-019-11809-2.

30. Worseck JM, Grossmann A, Weimann M, Hegele A, Stelzl U. A stringent yeast two-hybrid matrix screening approach for protein-protein interaction discovery. Methods Mol Biol. 2012;812:63–87. doi:10.1007/978-1-61779-455-1_4.

31. Manning G, Whyte DB, Martinez R, Hunter T, Sudarsanam S. The protein kinase complement of the human genome. Science. 2002;298:1912–34. doi:10.1126/science.1075762.

32. Casado P, Rodriguez-Prados J-C, Cosulich SC, Guichard S, Vanhaesebroeck B, Joel S, Cutillas PR. Kinase-substrate enrichment analysis provides insights into the heterogeneity of signaling pathway activation in leukemia cells. Sci Signal. 2013;6:rs6. doi:10.1126/scisignal.2003573.

33. Choteau SA, Cristianini M, Maldonado K, Drets L, Boujeant M, Brun C, et al. mimic INT: a workflow for microbe-host protein interaction inference; 2022.

34. Kumar M, Michael S, Alvarado-Valverde J, Mészáros B, Sámano-Sánchez H, Zeke A, et al. The Eukaryotic Linear Motif resource: 2022 release. Nucleic Acids Res. 2022;50:D497–D508. doi:10.1093/nar/gkab975.

35. Weatheritt RJ, Luck K, Petsalaki E, Davey NE, Gibson TJ. The identification of short linear motif-mediated interfaces within the human interactome. Bioinformatics. 2012;28:976–82. doi:10.1093/bioinformatics/bts072.

36. Parca L, Ariano B, Cabibbo A, Paoletti M, Tamburrini A, Palmeri A, et al. Kinome-wide identification of phosphorylation networks in eukaryotic proteomes. Bioinformatics. 2019;35:372–9. doi:10.1093/bioinformatics/bty545.

37. Zhang W, Zhang Y, Min Z, Mo J, Ju Z, Guan W, et al. COVID19db: a comprehensive database platform to discover potential drugs and targets of COVID-19 at whole transcriptomic scale. Nucleic Acids Res. 2022;50:D747–D757. doi:10.1093/nar/gkab850.

38. Kuleshov MV, Xie Z, London ABK, Yang J, Evangelista JE, Lachmann A, et al. KEA3: improved kinase enrichment analysis via data integration. Nucleic Acids Res. 2021;49:W304–W316. doi:10.1093/nar/gkab359.

39. Lonsdale J, Thomas J, et al. The Genotype-Tissue Expression (GTEx) project. Nat Genet. 2013;45:580–5. doi:10.1038/ng.2653.

40. Lachmann A, Torre D, Keenan AB, Jagodnik KM, Lee HJ, Wang L, et al. Massive mining of publicly available RNA-seq data from human and mouse. Nat Commun. 2018;9:1366. doi:10.1038/s41467-018-03751-6.

41. Subramanian A, Tamayo P, Mootha VK, Mukherjee S, Ebert BL, Gillette MA, et al. Gene set enrichment analysis: a knowledge-based approach for interpreting genome-wide expression profiles. Proc Natl Acad Sci U S A. 2005;102:15545–50. doi:10.1073/pnas.0506580102.

42. Subramanian A, Narayan R, Corsello SM, Peck DD, Natoli TE, Lu X, et al. A Next Generation Connectivity Map: L1000 Platform and the First 1,000,000 Profiles. Cell. 2017;171:1437–1452.e17. doi:10.1016/j.cell.2017.10.049.

43. Blanco-Melo D, Nilsson-Payant BE, Liu W-C, Uhl S, Hoagland D, Møller R, et al. Imbalanced Host Response to SARS-CoV-2 Drives Development of COVID-19. Cell. 2020;181:1036–1045.e9. doi:10.1016/j.cell.2020.04.026.

44. Dugourd A, Kuppe C, Sciacovelli M, Gjerga E, Gabor A, Emdal KB, et al. Causal integration of multi-omics data with prior knowledge to generate mechanistic hypotheses. Mol Syst Biol. 2021;17:e9730. doi:10.15252/msb.20209730.

45. Munk S, Refsgaard JC, Olsen JV, Jensen LJ. From Phosphosites to Kinases. Methods Mol Biol. 2016;1355:307–21. doi:10.1007/978-1-4939-3049-4_21.

46. Beekhof R, van Alphen C, Henneman AA, Knol JC, Pham TV, Rolfs F, et al. INKA, an integrative data analysis pipeline for phosphoproteomic inference of active kinases. Mol Syst Biol. 2019;15:e8981. doi:10.15252/msb.20198981.

47. Garrett S, Barton WA, Knights R, Jin P, Morgan DO, Fisher RP. Reciprocal activation by cyclin-dependent kinases 2 and 7 is directed by substrate specificity determinants outside the T loop. Mol Cell Biol. 2001;21:88–99. doi:10.1128/mcb.21.1.88-99.2001.

48. Fisher RP, Morgan DO. A novel cyclin associates with M015/CDK7 to form the CDK-activating kinase. Cell. 1994;78:713–24. doi:10.1016/0092-8674(94)90535-5.

49. Lee J-G, Huang W, Lee H, van de Leemput J, Kane MA, Han Z. Characterization of SARS-CoV-2 proteins reveals Orf6 pathogenicity, subcellular localization, host interactions and attenuation by Selinexor. Cell Biosci. 2021;11:58. doi:10.1186/s13578-021-00568-7.

50. Grossmann A, Benlasfer N, Birth P, Hegele A, Wachsmuth F, Apelt L, Stelzl U. Phospho-tyrosine dependent protein-protein interaction network. Mol Syst Biol. 2015;11:794. doi:10.15252/msb.20145968.

51. Corwin T, Woodsmith J, Apelt F, Fontaine J-F, Meierhofer D, Helmuth J, et al. Defining Human Tyrosine Kinase Phosphorylation Networks Using Yeast as an In Vivo Model Substrate. Cell Syst. 2017;5:128–139.e4. doi:10.1016/j.cels.2017.08.001.

52. Moesslacher CS, Kohlmayr JM, Stelzl U. Exploring absent protein function in yeast: assaying post translational modification and human genetic variation. Microb Cell. 2021;8:164–83. doi:10.15698/mic2021.08.756.

53. Wong HT, Cheung V, Salamango DJ. Decoupling SARS-CoV-2 ORF6 localization and interferon antagonism. J Cell Sci 2022. doi:10.1242/jcs.259666.

54. Yue M, Hu B, Li J, Chen R, Yuan Z, Xiao H, et al. Coronaviral ORF6 protein mediates inter-organelle contacts and modulates host cell lipid flux for virus production. EMBO J. 2023;42:e112542. doi:10.15252/embj.2022112542.

55. Miorin L, Kehrer T, Sanchez-Aparicio MT, Zhang K, Cohen P, Patel RS, et al. SARS-CoV-2 Orf6 hijacks Nup98 to block STAT nuclear import and antagonize interferon signaling. Proc Natl Acad Sci U S A. 2020;117:28344–54. doi:10.1073/pnas.2016650117.

56. Lei X, Dong X, Ma R, Wang W, Xiao X, Tian Z, et al. Activation and evasion of type I interferon responses by SARS-CoV-2. Nat Commun. 2020;11:3810. doi:10.1038/s41467-020-17665-9.

57. Miyamoto Y, Itoh Y, Suzuki T, Tanaka T, Sakai Y, Koido M, et al. SARS-CoV-2 ORF6 disrupts nucleocytoplasmic trafficking to advance viral replication. Commun Biol. 2022;5:483. doi:10.1038/s42003-022-03427-4.

58. Hall R, Guedán A, Yap MW, Young GR, Harvey R, Stoye JP, Bishop KN. SARS-CoV-2 ORF6 disrupts innate immune signalling by inhibiting cellular mRNA export. PLoS Pathog. 2022;18:e1010349. doi:10.1371/journal.ppat.1010349.

59. Lennartsson J, Ma H, Wardega P, Pelka K, Engström U, Hellberg C, Heldin C-H. The Fer tyrosine kinase is important for platelet-derived growth factor-BB-induced signal transducer and activator of transcription 3 (STAT3) protein phosphorylation, colony formation in soft agar, and tumor growth in vivo. J Biol Chem. 2013;288:15736–44. doi:10.1074/jbc.M113.476424.

60. Zoubeidi A, Rocha J, Zouanat FZ, Hamel L, Scarlata E, Aprikian AG, Chevalier S. The Fer tyrosine kinase cooperates with interleukin-6 to activate signal transducer and activator of transcription 3 and promote human prostate cancer cell growth. Mol Cancer Res. 2009;7:142–55. doi:10.1158/1541-7786.MCR-08-0117.

61. Dolgachev V, Panicker S, Balijepalli S, McCandless LK, Yin Y, Swamy S, et al. Electroporation-mediated delivery of FER gene enhances innate immune response and improves survival in a murine model of pneumonia. Gene Ther. 2018;25:359–75. doi:10.1038/s41434-018-0022-y.

62. Mizutani T, Fukushi S, Murakami M, Hirano T, Saijo M, Kurane I, Morikawa S. Tyrosine dephosphorylation of STAT3 in SARS coronavirus-infected Vero E6 cells. FEBS Lett. 2004;577:187–92. doi:10.1016/j.febslet.2004.10.005.

63. Tavares S, Liv N, Pasolli M, Opdam M, Rätze MAK, Saornil M, et al. FER regulates endosomal recycling and is a predictor for adjuvant taxane benefit in breast cancer. Cell Rep. 2022;39:110584. doi:10.1016/j.celrep.2022.110584.

64. Jackson CB, Farzan M, Chen B, Choe H. Mechanisms of SARS-CoV-2 entry into cells. Nat Rev Mol Cell Biol. 2022;23:3–20. doi:10.1038/s41580-021-00418-x.

65. V’kovski P, Kratzel A, Steiner S, Stalder H, Thiel V. Coronavirus biology and replication: implications for SARS-CoV-2. Nat Rev Microbiol. 2021;19:155–70. doi:10.1038/s41579-020-00468-6.

66. Pillaiyar T, Laufer S. Kinases as Potential Therapeutic Targets for Anti-coronaviral Therapy. J Med Chem. 2022;65:955–82. doi:10.1021/acs.jmedchem.1c00335.

67. Weisberg E, Parent A, Yang PL, Sattler M, Liu Q, Liu Q, et al. Repurposing of Kinase Inhibitors for Treatment of COVID-19. Pharm Res. 2020;37:167. doi:10.1007/s11095-020-02851-7.

68. Gutierrez-Chamorro L, Felip E, Ezeonwumelu IJ, Margelí M, Ballana E. Cyclin-dependent Kinases as Emerging Targets for Developing Novel Antiviral Therapeutics. Trends Microbiol. 2021;29:836–48. doi:10.1016/j.tim.2021.01.014.

69. Walhout AJ, Temple GF, Brasch MA, Hartley JL, Lorson MA, van den Heuvel S, Vidal M. GATEWAY recombinational cloning: application to the cloning of large numbers of open reading frames or ORFeomes. Meth Enzymol. 2000;328:575–92. doi:10.1016/s0076-6879(00)28419-x.

70. Dosztányi Z. Prediction of protein disorder based on IUPred. Protein Sci. 2018;27:331–40. doi:10.1002/pro.3334.

71. Hagai T, Azia A, Babu MM, Andino R. Use of host-like peptide motifs in viral proteins is a prevalent strategy in host-virus interactions. Cell Rep. 2014;7:1729–39. doi:10.1016/j.celrep.2014.04.052.

72. Love MI, Huber W, Anders S. Moderated estimation of fold change and dispersion for RNA-seq data with DESeq2. Genome Biol. 2014;15:550. doi:10.1186/s13059-014-0550-8.

73. Korotkevich G, Sukhov V, Budin N, Shpak B, Artyomov MN, Sergushichev A. Fast gene set enrichment analysis; 2016.

74. Wickham H. ggplot2: Elegant graphics for data analysis. Cham: Springer international publishing; 2016.

75. Shannon P, Markiel A, Ozier O, Baliga NS, Wang JT, Ramage D, et al. Cytoscape: a software environment for integrated models of biomolecular interaction networks. Genome Res. 2003;13:2498–504. doi:10.1101/gr.1239303.

76. Metz KS, Deoudes EM, Berginski ME, Jimenez-Ruiz I, Aksoy BA, Hammerbacher J, et al. Coral: Clear and Customizable Visualization of Human Kinome Data. Cell Syst. 2018;7:347–350.e1. doi:10.1016/j.cels.2018.07.001.

